# An optimized approach for multiplexing single-nuclear ATAC-seq using oligonucleotide conjugated antibodies

**DOI:** 10.1101/2022.12.22.521637

**Authors:** Betelehem Solomon Bera, Taylor V. Thompson, Eric Sosa, Hiroko Nomaru, David Reynolds, Robert A. Dubin, Shahina B. Maqbool, Deyou Zheng, Bernice E. Morrow, John M. Greally, Masako Suzuki

## Abstract

**Background:** Single-cell technologies to analyze transcription and chromatin structure have been widely used in many research areas to reveal the functions and molecular properties of cells at single-cell resolution. Sample multiplexing techniques are valuable when performing single-cell analysis, reducing technical variation and permitting cost efficiencies. Several commercially available methods are available and have been used in many scRNA-seq studies. On the other hand, while several methods have been published, the multiplexing techniques for single nuclear Assay for Transposase-Accessible Chromatin (snATAC)-seq assays remain under development. We developed a simple nucleus hashing method using oligonucleotide conjugated antibodies recognizing nuclear pore complex proteins, NuHash, to perform snATAC-seq library preparations by multiplexing.

**Results:** We performed multiplexing snATAC-seq analyses on the mixture of human and mouse cell samples (two samples, 2-plex, and four samples, 4-plex) using NuHash. The demultiplexing accuracy of NuHash was high, and only ten out of 9,144 nuclei (2-plex) and 150 of 12,208 nuclei (4-plex) had discordant classifications between NuHash demultiplexing and discrimination using reference genome alignments. We compared results between snATAC-seq and deeply sequenced bulk ATAC-seq on the same samples and found that most of the peaks detected in snATAC-seq were also detected in deeply sequenced bulk ATAC-seq. The bulk ATAC-seq signal intensity was positively correlated with the number of cell subtype clusters detected in snATAC-seq, but not the subset of peaks detected in all clusters. These subsets of snATAC-seq peaks showed different distributions over different genomic features, suggesting that the peak intensities of bulk ATAC-seq can be used to identify different types of functional loci.

**Conclusions:** Our multiplexing method using oligo-conjugated anti-nuclear pore complex proteins, NuHash, permits high accuracy demultiplexing of samples. The NuHash protocol is straightforward, it works on frozen samples, and requires no modifications for snATAC-seq library preparation.

## Introduction

Advancing single-cell technologies to analyze chromatin structure and transcription profiles allows us to assess the transcription regulatory and transcriptomic landscapes of each cell subtype within a heterogeneous sample. The single-nuclear ATAC-seq (snATAC-seq) assay is based on the <u>a</u>ssay for <u>t</u>ransposase-<u>a</u>ccessible <u>c</u>hromatin (ATAC) to define sites of open chromatin in the genome [1, 2], thus identifying regulatory loci at a single-cell resolution. When we assess these regulatory landscapes at single cell resolution, a single nucleus is captured in an oil droplet containing a barcoded capture bead or sorted into a single well, and then a library is generated for each nucleus in an isolated environment. These steps are usually performed on a sample-by-sample basis, potentially resulting in technical batch effects on the results that are sometimes hard to resolve computationally at the analysis step. To address this technical difficulty, several multiplexing methods have been developed and widely used to reduce technical batch effects in single-cell RNA-seq (scRNA-seq) studies [3–8]. Recently, Zhang *et al.* reviewed the characteristics of sample-multiplexing approaches used for single-cell sequencing [9]. Among those, the methods using natural genetic variation, such as single nucleotide variants (SNVs), do not require a step prior to generating a scRNA-seq library [6, 10–12]. However, genetic variation-based methods are not applicable to studies lacking genetic differences between samples, such as model organism studies using congenic strains. The most accepted method is cell hashing, defined as pooling sets of cells, utilizing uniquely barcoded oligo-conjugated antibodies [4] or lipids to tag each of the samples [7]. The first step for both methods is to incubate the cells with an antibody or lipids that binds to an epitope present on the cells being tested in a given sample. Individual samples are labelled with differently barcoded antibodies, and then the samples are combined for a single assay. The conjugated oligonucleotides with a unique barcode will subsequently be sequenced in the single cell assay, and the unique barcode is utilized to demultiplex the combined samples. This has the effect of minimizing the technical batch effects that would otherwise make assays difficult to compare and reduces the amount of sequencing needed. Barcoded oligonucleotides conjugated antibodies and lipids are commercially available and widely used in scRNA-seq analysis for demultiplexing. However, since these conjugated oligonucleotides are designed to be captured by Oligo-dT scRNA-seq probes, these antibodies or lipids are not applicable to snATAC-seq. There currently are only a few multiplexing techniques available for snATAC-seq [8, 13–15]. These techniques are complicated and sometimes increase the number of steps needed to generate barcoded libraries or require modifying the library preparation method.

In this study, we developed a simple <u>nu</u>cleus <u>hash</u>ing method, NuHash, to perform snATAC-seq library preparations by multiplexing, a similar procedure to the cell hashing method used for scRNA-seq. We designed hashing oligonucleotides containing a Tn5 tag sequence and a specific barcode that can be sequenced in the single-cell assay, with the oligonucleotides conjugated to an antibody to a nuclear membrane protein complex. Using NuHash, we demonstrated the capabilities of multiplexing samples in both humans and mice. In addition, our analysis revealed the importance of evaluating peak intensities when interpreting data from bulk ATAC-seq.

## RESULTS

### Design overview

A flow diagram of our library preparation method at each step is shown in **Figure 1**. The left side shows the ATAC-seq products, and the right side the NuHash products. The NuHash oligonucleotide was conjugated to the anti-Nuclear Pore Complex Proteins antibody. The NuHash oligonucleotide contains two unique molecular identifier sequences (UMI), the sample hashing sequence located between the UMIs, a part of Illumina Read 1 sequence at the 5’ end, and a part of Illumina Read 2 sequence is on the 3’end. We included two phosphorothioate bonds at the 3’ end of the sequence to protect the oligonucleotide-probes from cell nucleases (**Supple Table 1**). The Illumina Read1 sequence part binds to the gel beads of during the Gel Beads-in-emulsion (GEM) step, and it acquires the complete sequence combination after amplification of the library.

**Figure 1:**
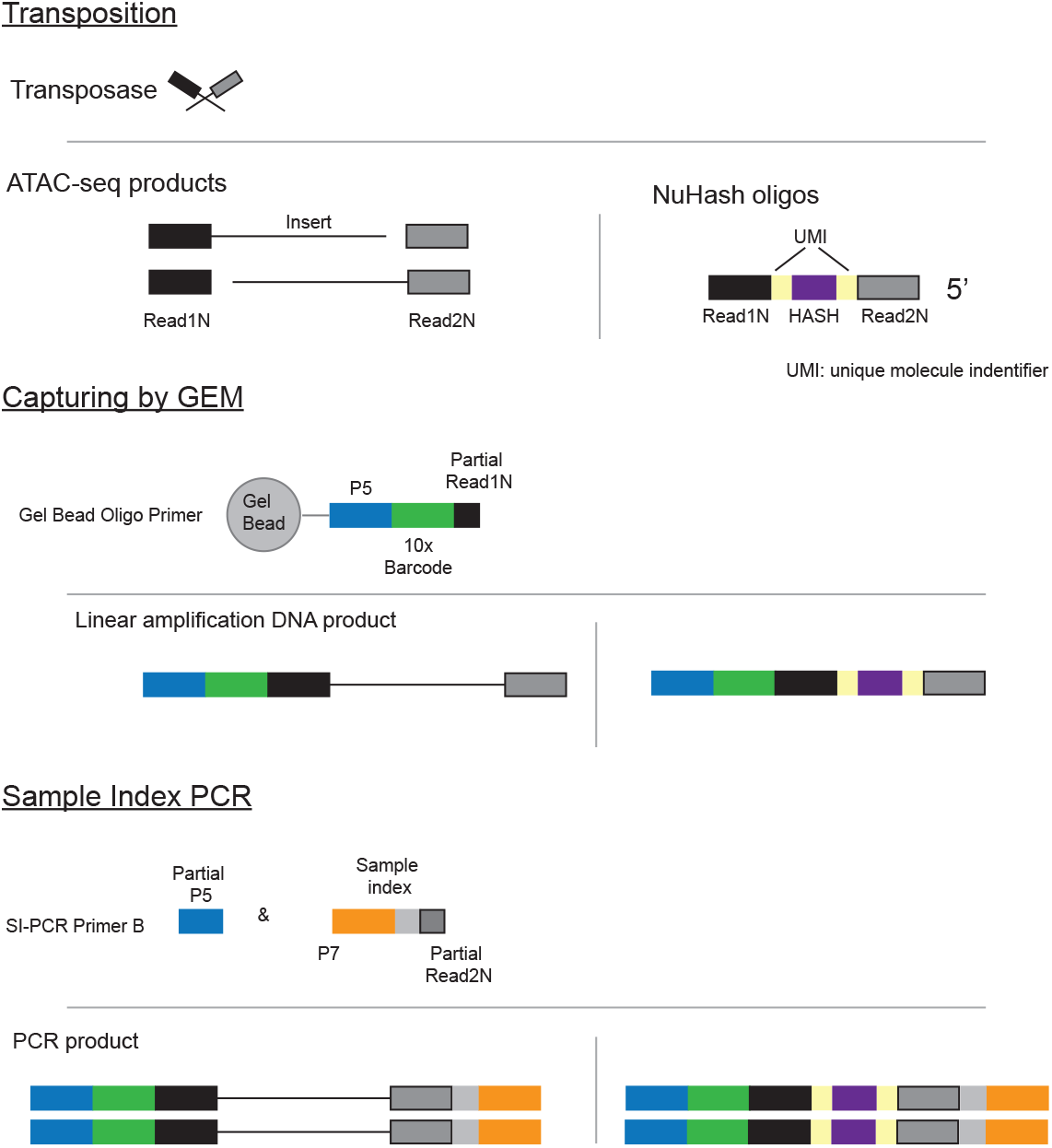
Overview of library preparation of NuHash. We illustrated the product’s progression step by step. The right panel shows the NuHash library production, and the left shows the ATAC-seq library production.

We performed two independent library preparations on mixtures of human and mouse nuclei to assess the accuracy of demultiplexing by alignment results. We used frozen human CD4+ T cells and mouse hematopoietic progenitor cells line (HPC-7), and each sample was stained with a different NuHash antibody. For the first set, we used two samples; one human CD4+ T sample and one HPC-7 sample. For the second set, we used four samples in total, two samples of CD4+ T cells (male and female, allowing us to assess the accuracy of demultiplexing by alignment results) and two samples of HPC-7. Hereafter, we call the first set 2-plex and the second 4-plex. After staining with different NuHash antibodies, we evenly combined samples of nuclei and adjusted to 7,086 nuclei/sample/μl (2-plex) or 7,340 nuclei/sampe/μl (4-plex) to target 10,000 and 20,000 nuclei per snATAC-seq experiment. We assessed the quality of the libraries before sequencing by examining the fragment analyzer traces, observing the expected banding pattern of the fragment distributions [1, 2] (**Supple Figure 2**).

### Library sequencing and assessing qualities of the NuHash snATAC-seq libraries

We sequenced the libraries on the Illumina NextSeq 500 sequencer (50 bp for read1, 8 bp for i7 index read, 16 bp for i5 index read, and 50 bp for read 2). The paired-end sequence reads were aligned to a human (GRCh38) and mouse (mm10) combined reference genome using Cell Ranger ATAC software (version 2.0.0). The total numbers of read pairs and detected cells were 446,720,359 and 16,262 for 2-plex, respectively, and 353,484,777 and 34,248 for 4-plex, respectively. The alignment statistics were summarized in **Supple Table 2**. We assessed the quality of the snATAC-seq libraries using ArchR [16]. From the ArchR analysis, we detected 2,266 and 12,880 nuclei aligned to the human genome in 2-plex and 4-plex, respectively. The median fragment numbers per nuclei were 2,103 (2-plex) and 1,885 (4-plex), and the median transcription start site (TSS) enrichment were 22.69 (2-plex) and 22.808 (4-plex), respectively (**Supple Figure 3A**). The detected duplex was 51 of 2,266 (2.3%, 2-plex) and 1,658 of 12,880 (12.9%, 4-plex), respectively (**Supple Figure 3B**). In mouse alignment data, we detected 4,360 and 13,054 nuclei aligned to the mouse genome in 2-plex and 4-plex, respectively. The median fragment numbers per nuclei were 2,731 (2-plex) and 2,280 (4-plex), and the median TSS enrichments were 21.09 (2-plex) and 22.69 (4-plex), respectively (**Supple Figure 3C**). The detected duplex was 0 of 4,360 (0%, 2-plex) and 1,704 (13.1%, 4-plex), respectively (**Supple Figure 3D**). We observed the expected banding patterns in the insert fragment length distribution in all libraries (**Supple Figure 3E**).

### Accuracy of NuHash

We then selected nuclei that have at least 10,000 read counts for further analysis, resulting in 10,574 nuclei (2-plex) and 12,208 nuclei (4-plex) using Signac [17]. We counted the number of hashing sequences per nucleus. We plotted the human and mouse genome aligned read number per nucleus (**Figure 2A**) and the number of hash sequence counts for Oligo 1 (human) and Oligo 2 (mouse) (**Figure 2B**) of the 2-plex data. We observed a clear dissociation between human and mouse nuclei based on the alignment status and the hash sequence counts. In **Figure 2C**, we plotted the number of reads aligned to mouse or human reference and colored based on NuHash sequence status. We assessed the nuclear capture by testing the alignment rates of the reference sequence and the counts of NuHash per nucleus. We classified nuclei as singlet if the reads aligned to one of the reference genomes and doublet if the reads aligned to both. For NuHash, we classified nuclei as singlet if the NuHash reads aligned to a single NuHash barcode sequence, and duplicate if the NuHash reads aligned to two or more different NuHash barcode sequences. We detected 9,144 (86.48%) singlet nuclei and 1,260 (11.92%) duplicate nuclei, while 170 (1.61%) nuclei did not have enough NuHash counts (NA). Of the 9,144 singlet nuclei, 4,409 (human/oligo 1) and 4,735 (mouse/oligo 2) were classified based on the NuHash count status. We compared the nucleus assigned classification (human singlet, mouse singlet, and duplicates) based on genome read alignments and NuHash count status, and detected only 10 discordantly classified nuclei (8 human/oligo2 and 2 mouse/oligo 1, p-value < 2.2×10^-16^, Chi-squared test). The calling accuracy on the 4-plex experiment was comparable to the 2-plex experiment (p-value < 2.2×10^-16^, Chi-squared test). We detected 12,208 singlets of those 7,079 human nuclei with accurate NuHash information, 4,979 mouse nuclei with accurate NuHash information, 974 newly classified, and only 150 nuclei with the discordant classifications (**Supple Figure 4**).

**Figure 2:**
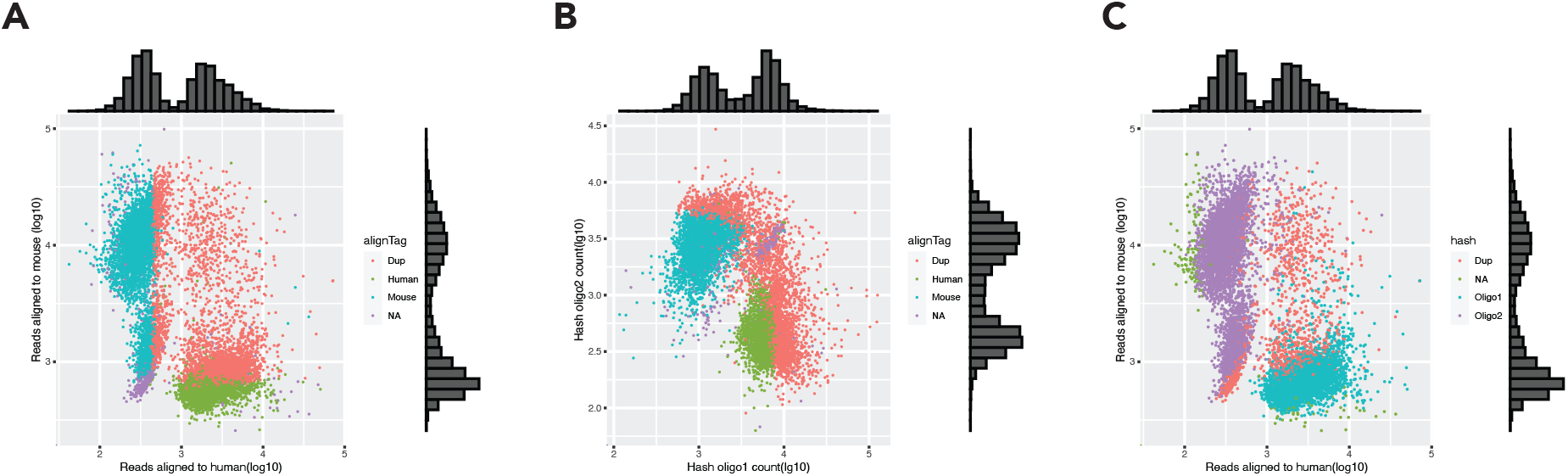
The fragment alignment status of each nucleus. The fragment alignment status of each nucleus is shown in three panels. (**A**) The number of read saligned to human or mouse references per nucleus, (**B**) the NuHash sequence counts per nucleus, and (**C**) the number of reads aligned to human or mouse references per nucleus colored by the NuHash demultiplexed status.

### Clustering analysis

We performed single-cell clustering analysis to identify clusters of cells on human and mouse genome-aligned nuclei. We eliminated the nuclei if one of the following conditions was met; 1) the percent of reads in peaks was less than 15%, 2) the ratio of reads aligned to the genomic blacklist (loci with anomalous, unstructured, or high signal in next-generation sequencing experiments independent of cell line or experiment [18]) was >0.05, 3) nucleosome signal was <0.2 or >4, and 4) the number of fragments aligned to peaks was <2,000. A total of 2,920 human nuclei and 3,311 mouse nuclei passed these thresholds. Among those, we detected 5 clusters in human and 5 clusters in mouse (**Figure 3A top**). As we expected, very few hashed mouse nuclei were clustered in the human nuclei cluster, and *vice versa* (**Figure 3A bottom, Supple Figure 5**). We plotted the aligned reads for the human *EIF1AY* gene that is located on chromosome Y as a representation of sample hashing accuracy. We observed an enrichment only in the Human 2 (male) sample (**Figure 3B**). This result supports the high demultiplexing accuracy of NuHash.

**Figure 3:**
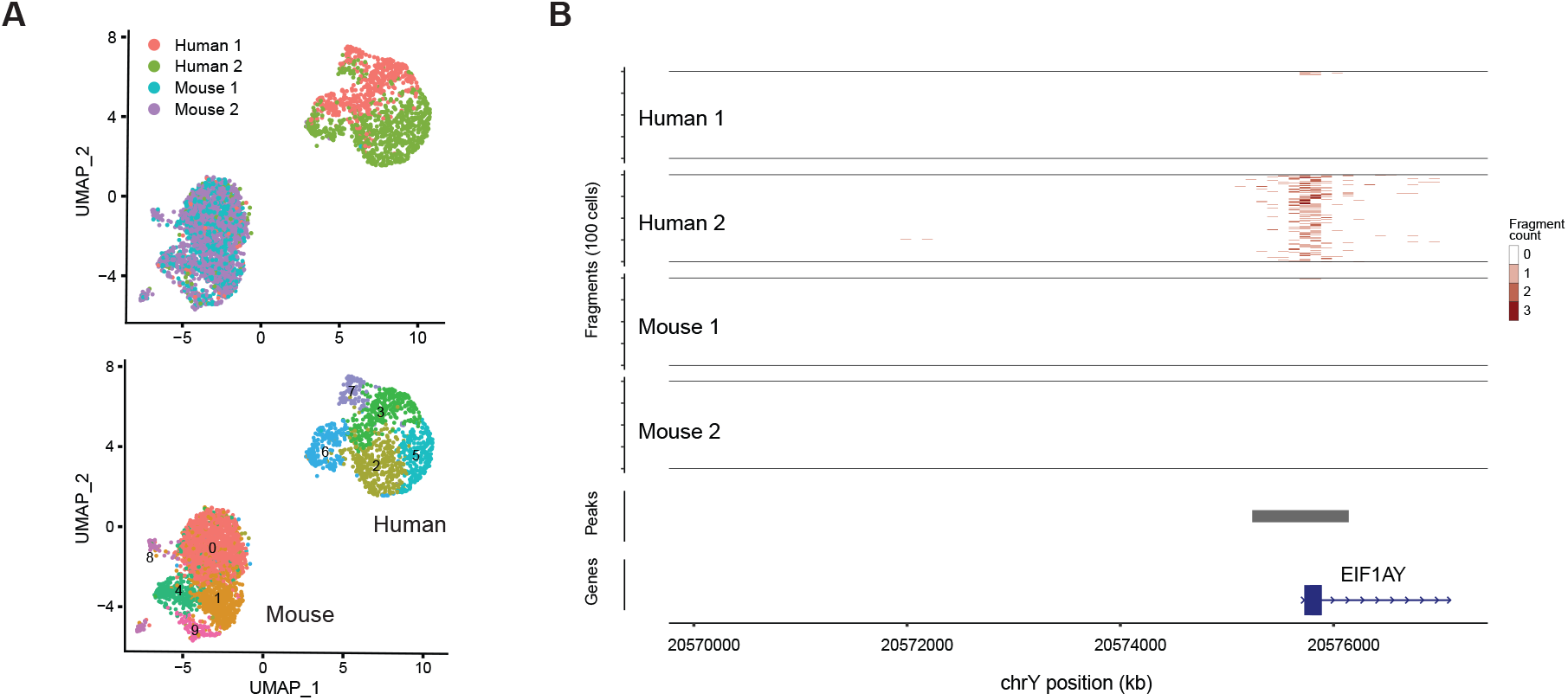
The snATAC-seq analysis identified distinct cell clusters by their open chromatin status. The clustering analysis based on open chromatin status identified ten cell clusters (five mouse and five human cell clusters). (**A**) Identified cell clusters were visualized by UMAP, and each nucleus was colored by the cell clusters (top) or NuHash demultiplexed status (bottom). (**B**) A representative view of sample-specific reads alignment at the promoter region. At the promoter region of the human *EIF1AY* gene, only reads of the human 2 (male) sample were detected in this chromosome Y region.

### Bulk ATAC-seq

To assess the accuracy of our NuHash approach, we performed an ultra-deep bulk ATAC-seq on HPC7 mouse cells to obtain a well-defined open-chromatin region (OCR) profile of the cell line. A total of 8 ATAC-seq libraries from independent cell culture batches were sequenced with the goal to obtain 50 million paired-end reads per sample. We excluded one sample which did not have more than 10,000 paired reads. The sequencing statistics of each library are summarized on **Supple Table 3**. We obtained a median of 69.8 million paired-end reads per sample (a total 700.7 million paired-end reads used in the analysis), with the median percentage of reads in OCRs of 58.8%, the median percent of duplication 0.22%, and the detected OCRs ranging from 84,593 to 13,2852 (mean=105,983, standard deviation =17,024). First, we assessed the reproducibility of the OCRs between libraries. After intersecting OCRs from seven libraries, we identified 92,740 intersected OCRs, of which 11,058 were significantly overlapped in at least 3 libraries (high-confident OCRs, hcOCRs). As we expected, the hcOCRs are enriched near TSS (TSS to 1 kb upstream, 58.9%) (**Supple Figure 6A**). The motif analysis on the hcOCRs showed enrichment of hematopoietic lineage-related transcription factor genes, such as *Elk1, Runx2,* and *Klf7* (**Supple Figure 6B**). We then assessed the overlap of the hcOCRs with snATAC-seq detected peaks. Of the 11,508 hcOCRs, only 46 peaks were not detected in snATAC-seq peaks, suggesting most of hcOCRs are detected in data from NuHash snATAC-seq experiments.

### Cell subtype peaks

We assessed the characteristics of snATAC-seq peaks by categorizing them by the number of cell subtype clusters (clusters) in which the peak was present. To increase the robustness of the analysis, we re-performed clustering only for mouse nuclei and selected clusters containing at least 50 nuclei (**Figure 4A**). We identified 85,951 peaks and four clusters in mouse HPC7 snATAC-seq data, of which 77,987 peaks overlapped with OCRs detected in bulk ATAC-seq (see previous section). Of these 77,987 peaks, 16,810 peaks were detected in only one cluster (Cnum_1), 11,964 in two clusters (Cnum_2), 12,743 in three clusters (Cnum_3), and 36,470 in all four clusters (Cnum_4). When we then looked at the peak intensity of the overlapped bulk ATAC-seq OCRs, they were positively correlated with the number of clusters with peaks (**Figure 4B**). The constitutive Cnum_4 peaks were enriched in promoters and 5’UTR regions (**Figure 4C**). Interestingly, the peak heights of the Cnum_4 bulk ATAC-seq OCRs were bimodally distributed, suggesting the existence of stochastic OCRs (low-intensity peaks) and constant OCRs (high-intensity peaks) (**Figure 4B**). The high-intensity peaks were enriched in promoters and 5’UTR regions, while the low-intensity peaks were enriched in gene bodies. This finding suggests that molecular mechanisms that create constitutively open chromatin differ between the stochastic and constant OCRs.

**Figure 4:**
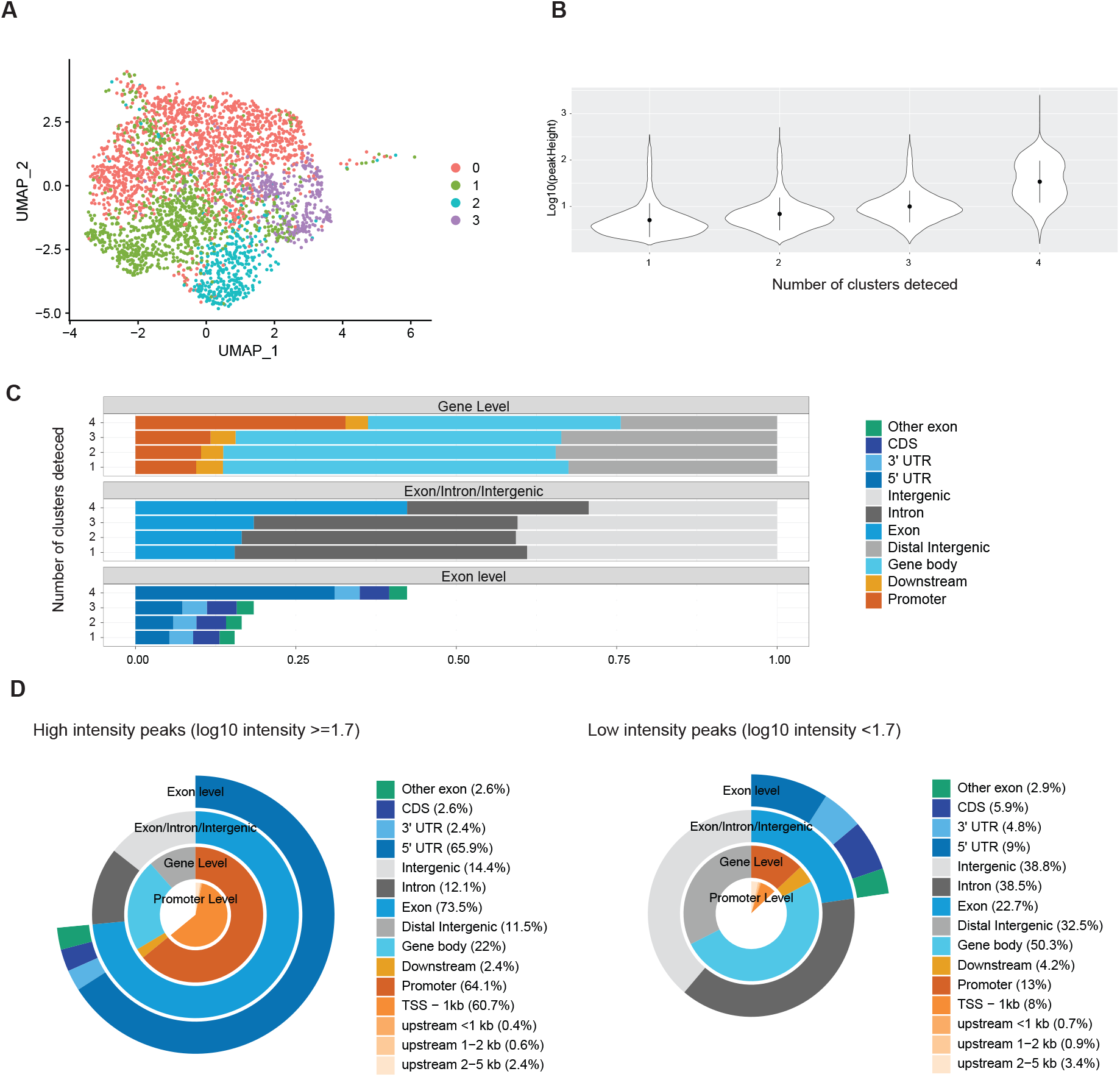
An integration analysis of snATAC-seq and deeply sequenced bulk ATAC-seq. (**A**) a UMAP plot shows the mouse cell clusters identified in the snATAC-seq analysis, (**B**) the peak intensities of overlapped bulk ATAC-seq peaks were plotted by the number of detected cell clusters in snATAC-seq, (**C**) distributions over different genomic features of the peaks categorized by the number of detected cell clusters, and (**D**) distributions over different genomic features of the peaks detected in all clusters dichotomized by the peak intensities of overlapped bulk ATAC-seq peaks.

We also assessed the gene expression properties for each peak category group. As expected from the location of the OCRs, the genes containing Cnum_4 high-intensity peak group have higher expression levels compared to other groups (**Figure 5A**). We also assessed overlap of each group to transcriptionally active putative enhancers with potential bidirectional transcription (TAPEs) [19]. We identified 145 TAPEs in HPC7 cells using our deeply sequenced ATAC-seq data and publicly available HPC7 RNA-seq data (**Supple Table 4**). Of those, 92 TAPEs were also detected in the snATAC-seq data. More than 70% TAPEs identified from the snATAC-seq data overlapped with Cnum_4 peaks. Interestingly, they overlapped in both low- and high-intensity groups (**Figure 5B**). When we intersected Cnum_4 peaks with the reported transcription factor ChIP-seq peaks, hematopoietic lineage-related transcription factors were enriched in the high-intensity peak group (**Figure 5C**). Of note, a comparison of motif enrichment analysis results between low- and high-intensity groups revealed that the CTCF/BORIS motif was significantly enriched in the low-intensity peak group (**Figure 5D**). We further investigated the characteristic differences between low and high intensity peaks overlapping with the CTCF ChIP-seq peaks. The proportions of overlap with CTCF ChIP-seq peaks (CTCF peaks) were significantly higher in Cnum_4 groups (low-intensity, 41.9%, chi-square statistic =4038.4864, p < 0.00001; high-intensity, 40.2%, chi-square statistic=2105.6447. p < 0.00001) than all peaks detected in bulk ATAC-seq (22.14%) (**Figure 5E**). The absolute distances of the CTCF peaks from the TSSs were significantly shorter in the high-intensity peak group as compared to the low-intensity peak group or all bulk ATAC-seq peaks (**Figure 5F**). Since CTCF and cohesins are master regulators of topologically associating domains (TADs), we also tested if these CTCF peaks were located in promoter-interacting regions (PIRs) [20, 21]. While only 12.7% of CTCF peaks in low-intensity groups overlapped with PIRs, 60.4% of CTCF peaks in high-intensity groups overlapped with PIRs, suggesting functional CTCF peaks might be enriched in high-intensity groups.

**Figure 5:**
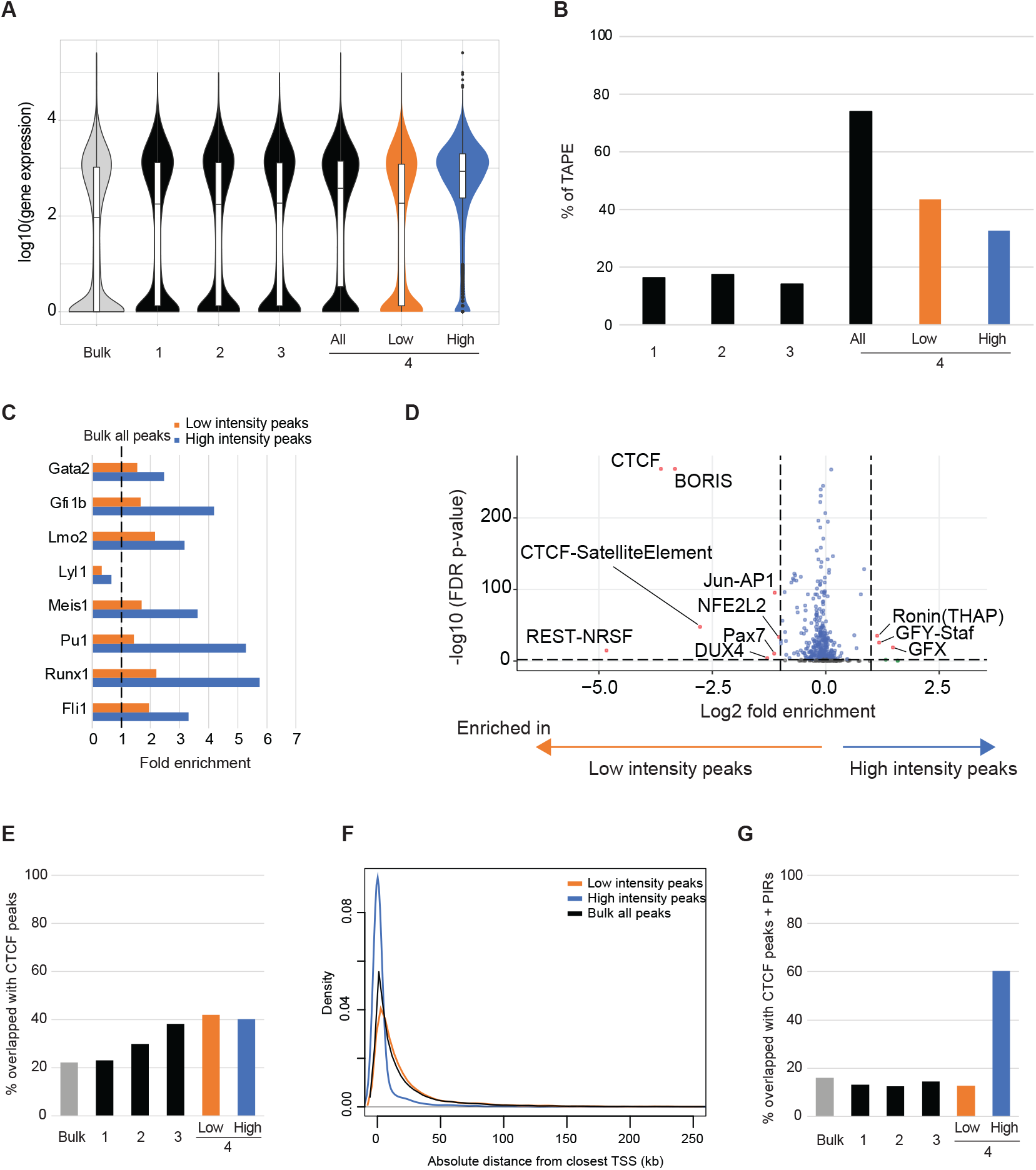
Differences in peak characteristics by the number of clusters detected. (**A**) Expression status of the genes located near the peaks, (B) the proportions of peaks overlapped with TAPEs, (C) comparisons of hematopoietic transcription factor binding motif enrichment of Cnum_4 peaks compared to all bulk ATAC-seq peaks, (D) a comparison of transcription factor binding motif enrichment between low- and high-intensity Cnum_4 peaks, (E) % overlapped with CTCF ChIP-seq peaks, (F) the distribution of absolute distance from TSS of Cnum_4 peaks, and (G) % overlapped with CTCF ChIP-seq peaks and PIRs.

## Discussion

In this study, we developed a simple nucleus hashing method, NuHash, to perform multiplexing using a high-throughput droplet-based snATAC-seq platform *(e.g.,* 10X Genomics), that is based upon a similar principle to the Cell Hashing technique for scRNA-seq multiplexing [4]. We tested the assay of two (2-plex) or four (4-plex) human and mouse samples to assess the capabilities of NuHash. We detected 9,144 and 12,208 tagged single-nuclei, respectively. Both 2-plex and 4-plex analyses showed high accuracy of demultiplexing samples using the barcode sequence of the conjugated oligonucleotide; only 0.11% and 1.23% of nuclei showed mismatched calling between hashing results and aligned DNA results, respectively. This result indicated that our simple nucleus hashing approach, NuHash, can effectively demultiplex the combined samples in snATAC-seq analysis.

The results allowed some insights into the properties of the signals observed. Comparisons between snATAC-seq and deeply sequenced bulk ATAC-seq of mouse hematopoietic progenitor cells revealed the importance of studying peak intensities of ATAC-seq data in analyses. As expected, the deeply sequenced bulk ATAC-seq peak intensities were positively correlated with the number of cell clusters detected, i.e., lower intensity peaks were called in a subset of cell clusters. However, some low-intensity peaks were detected in all cell subtypes (Cnum_4 peaks), resulting in a bimodal distribution of bulk ATAC-seq peak intensity in the Cnum_4 peaks. This result suggests the existence of metastable OCRs in the mouse genome. We further assessed the peak characteristic differences between low- and high-intensity Cnum_4 peaks by integrating with RNA-seq and transcription factor ChIP-seq data, revealing that higher intensity Cnum_4 peaks were detected at highly expressed genes, and that hematopoiesis-related transcription factor binding sites were enriched in higher intensity Cnum_4 peaks. In contrast, although snATAC-seq detected TAPEs were enriched in Cnum_4 peaks, the rate of overlap of TAPEs was comparable in lower- and higher-intensity peaks. A transcription factor motif enrichment analysis showed that CTCF-binding motifs were enriched in low-intensity Cnum_4 peaks. Moreover, CTCF-binding loci identified in ChIP-seq data were comparably enriched in both low- and high-intensity Cnum_4 peaks. However, as we expected from the peak annotation results, low-intensity CTCF peaks were located far from the TSS, and high-intensity CTCF peaks were closer to the TSS. In addition, these high-intensity CTCF peaks were enriched in the PIRs; thus, these high-intensity peaks may identify the transcription regulatory regions within the boundaries of TADs. This finding suggests that even though the peaks overlapped with CTCF peaks, low-intensity peaks are less likely to contribute to the regulatory functions in gene expression, and instead might be located in metastable binding regions (**Figure 6**).

**Figure 6:**
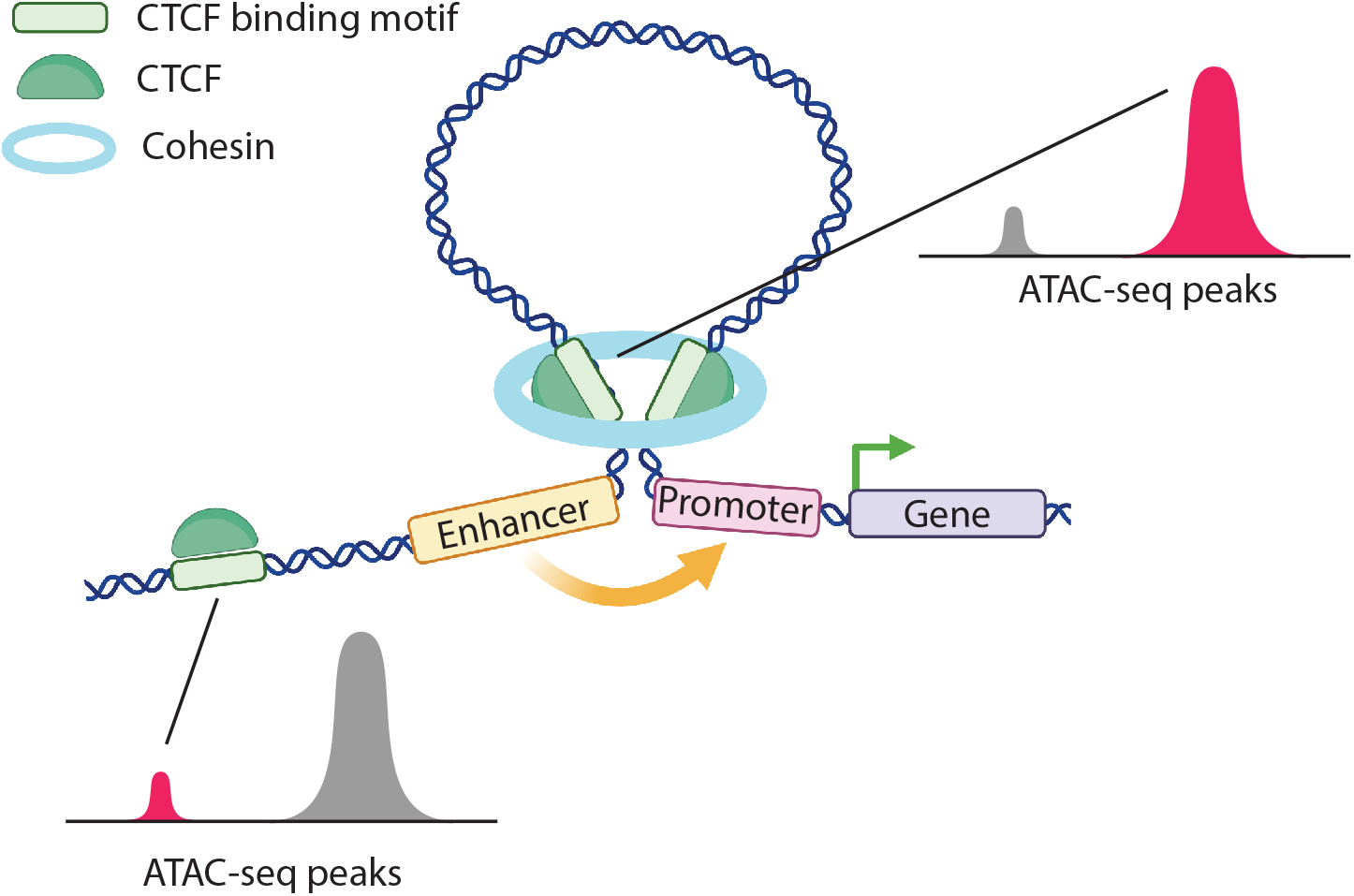
Proposed model of low- and high-intensity peaks and their associations to transcriptional regulations. Our results suggested that we could dissociate the functional enhancers by the existence of CTCF binding motifs and ATAC-seq peak intensities. Adapted from “Transcriptional Regulation by CTCF and Cohesin”, by BioRender.com (2022, https://app.biorender.com/biorender-templates)

To date, there are several multiplexing techniques that have been described to perform a high-throughput droplet-based snATAC-seq by multiplexing, such as dsciATAC-seq [14], CASB [8], and SNuBar [15]. The dsciATAC-seq method uses indexed Tn5 transposon complexes to tag the samples [14], CASB tags the samples using concanavalin A with biotinylated oligonucleotides and streptavidin [8], and SnuBar adds barcoded oligonucleotides at the tagmentation step before partitioning on a microfluidic chip [15]. Our NuHash method uses a stable oligoconjugated antibody for indexing and works on frozen samples. Both dsciATAC-seq and SnuBar tag the samples at the Tn5 tagmentation step, while CASB and NuHash tag the samples before the tagmentation step. While dsciATAC-seq requires a customized Tn5 enzyme, the other three do not. Since CASB and NuHash tag the samples before the actual snATAC-seq procedure, no modifications are required during the snATAC-seq library preparation. NuHash uses oligonucleotide-conjugated antibodies, removing any complicated optimization for tagging.

While the current snATAC-seq technique is designed to generate a library from up to 10,000 nuclei, we speculate that the analyzable nucleus number of snATAC-seq will be increased by advancing technology in the near future, given how the latest scRNA-seq method can analyze up to 3,500,000 nuclei with multiplexing. We believe snATAC-seq with multiplexing should become a common technique in the future. As a side note, the antibody used in this study has reactivities with nuclear pore complex proteins of vertebrates, *Xenopus,* and yeast; therefore, this hashing method could be used in the assays for other species, as well as in multi-omics studies which use isolated nuclei. There are some limitations of our NuHash: for instance, performing multiplexing analysis requires another 10 to 15% of sequence reads for demultiplexing. Also, titrating antibodies using the same sample type (species, tissue, and cell types) might be required before performing a large set of samples to obtain high-quality libraries. However, these limitations are not unique to NuHash, as most other multiplexing techniques have the similar limitations.

## Conclusion

We have developed a new simple method, NuHash, to perform snATAC-seq analysis with multiplexing using oligo-conjugated anti-nuclear pore complex proteins, which can be used for frozen samples, and demonstrated the accuracy of demultiplexing of NuHash. An integration analysis of snATAC-seq with NuHash and deeply sequenced bulk ATAC-seq datasets revealed the importance of considering peak intensity in interpreting the bulk ATAC-seq results.

## Materials and Methods

Detailed NuHash protocol is provided in the Supplementary information document.

### Custom oligo conjugated nucleus hashing antibody

The sequences of the custom oligo are listed in **Supple Table 1**. The custom oligos were synthesized by Integrated DNA Technologies and conjugated to anti-Nuclear Pore Complex Proteins antibody (Clone Mab414, BioLegend) by BioLegend. The oligo conjugated antibodies were aliquoted and stored at 4 °C until use.

### CD4 T-cell isolation from whole blood

CD4 T-cells were isolated from 10-20 ml of peripheral blood using EasySep Direct Human CD4+ T cell Isolation kit (Stem Cell Technologies, cat #19662). The isolated CD4 T-cells were stored in Crystor CS10 (Stem Cell Technologies, cat #07930) at −80 °C until use (50,000 cells per tube). This study was approved by the Albert Einstein College of Medicine Institutional Review Board (IRB Protocol# 2021-12969 and 2007-272).

### HPC-7 Hematopoietic progenitor cell

Hematopoietic progenitor cell line HPC-7 was kindly gifted by Dr. Britta Will at Albert Einstein College of Medicine. HPC-7 cells were maintained at density of 2-10 x10^5^/ml in Iscove’s modified Dulbecco’s medium (Invitrogen) supplemented with 50 ng/ml of mouse stem cell factor (Gemini Bio-Products), 1 mM Sodium Pyruvate, 6.9 ng/mL α-Monothioglycerol (SIGMA-Aldrich), 5% of bovine calf serum and Penicillin-Streptomycin.

### Nucleus isolation

For frozen human CD4 T cells were thawed with a series of dilutions with pre-warmed 10% FBS (Gemini Bio, Cat# 100-106) supplemented RPMI-1640 medium (Gibco, cat# 11875093), and washed with 0.04% BSA (Sigma-Aldrich, cat# 126609-5GM) in PBS(-) (Gibco, cat# 10010031). Freshly collected HPC-7 cells were washed with 0.04% BSA in PBS(-) before the lysis. We lysed pelleted cells with lysis buffer (10 mM Tris-HCl (pH 7.4), 10 mM NaCl, 3 mM MgCl2, 0.1% Tween-20, 0.1% NP-40, 0.01% Digitonin (Invitrogen, cat# BN2006)) for 3 minutes on ice, then we washed nuclei with wash buffer (10 mM Tris-HCl (pH 7.4), 10 mM NaCl, 3 mM MgCl2, 1% BSA, 0.1% Tween-20) and followed by washing with staining buffer (2% BSA, 0.01% Tween-20 in PBS(-)). The isolated nuclei were resuspended in 10 μl of the staining buffer. The number of nuclei in the solution was counted using hemocytometer (Fisher Scientific, cat# 0267151B).

### Nucleus hashing antibody staining

We stained isolated nuclei with nucleus hashing antibody in the staining buffer. After counting the number of nuclei per μl of nuclei suspension, we adjusted the number of nuclei 150,000 −500,000 in 100 μl of the staining buffer. After the Fc blocking with 10 μl of FcX (BioLegend, cat# 422301/ 101319) for 10 minutes on ice, we stained nuclei with nucleus hashing antibody for 20 minutes on ice, then washed the nuclei three times with 1 ml of the staining buffer. After the last wash step, we removed all supernatant and resuspended the nuclei with 5 μl of Diluted Nuclei buffer (10x Genomics, PN-20000153/20000207). The number of nuclei in the solution was counted using hemocytometer (Fisher Scientific, cat# 0267151B).

### Chromatin accessibility assay

Chromatin accessibility (ATAC-seq) assay was performed according to the Omni-ATAC protocol with some modifications [2]. Freshly isolated nuclei were spun down (500 RCF, 10 min, 4 °C), the supernatant was carefully removed, and nuclei pellet was resuspended in 50 μL of the transposase reaction mix including 25 μl of 2xTD buffer (Illumina, cat#15027866), 2.5 μl of transposase (Illumina, cat#15027865), 16.5 μl of PBS(-), 0.5μl of 1% digitonin (Promega, cat# G9441), 0.5 μl of 10% Tween-20, 5 μl of nuclease free H2O. The transposition reaction was performed at 37 °C for 30 minutes, followed by purification using Zymo DNA Clean and Concentrator-5 kit (Zymo Research, cat# D4013). A purified, transposed DNA was eluted in 11 μL of EB elution buffer and stored at *-*20 °C until amplification. For indexing and amplification of transposed DNA, we combined the following for each sample: 10 μL of transposed DNA, 25 μl of NEBNext High-Fidelity 2x PCR Master Mix (New England Biolabs, M0541S), 2.5 μL each of Nextera i5 and i7 indexed amplification primers (Nextera Index Kit, Illumina, FC-121—1011) and 10 μl of nuclease free H2O. The PCR reaction was carried out using the following conditions: one cycle of 72 °C for 5 minutes and 98 °C for 30 seconds; followed by ten cycles of 98 °C for 10 seconds, 63 °C for 30 seconds and 72 °C for 1 minute; and a hold step at 4 °C. The libraries were purified with a double-sided beads purification using AMPure XP (Beckman Coulter, catalog # A63880) and eluted in 20 μL of the elution buffer. The library quality was assessed by Bioanalyzer High-Sensitivity DNA Assay. The ATAC-seq libraries were quantified by Qubit HS DNA kit (Life Technologies, Q32851). 150 bp, paired-end sequencing was performed on a HiSeq 2500 Illumina instrument at the Novogene Co., Ltd.

### Antibody titration analysis

We performed antibody titration assays to obtain optimal concentrations for the nucleus-hashing antibody. We stained isolated 50,000 human CD4 T-cell nuclei at the ratios of 0.01 μg of nucleus-hashing antibody per 10000, 25000, and 50000 nuclei. After staining, the nuclei were washed three times with staining buffer, then the libraries were generated using Omni-ATAC protocol [2]. The ratios of nucleus-hashing products and the ATAC-seq products were assessed by Bioanalyzer High-Sensitivity DNA Assay.

### Single-nuclei ATAC-seq library preparation

The single-nuclei ATAC-seq libraries were generated using Chromium Single Cell ATAC seq library preparation kit (10x Genomics, cat# PN-1000111/ PN-1000084). The nuclei stained with nucleus-hashing antibodies were adjusted at the concentration of 7000 nuclei/μl or 7700 nuclei/μl and combined the two or four samples stained with different antibodies into a tube to run the library preparation by following the manufacturer’s instructions. The library preparation step was performed at the Genomics Core at Albert Einstein College of Medicine. After amplification, we sequenced the libraries as follows; 50 bp for read1, 8 bp for i7 index read, 16 bp for i5 index read, and 50 bp for read 2. Sequencings were performed at the Epigenomic Shared Facility at Albert Einstein College of Medicine.

### Nucleus-hashing analysis

The sequence reads were aligned to a reference genome that combined the human (GRCh38) and the mouse (mm10) reference genomes (refdata-cellranger-atac-GRCh38-and-mm10-1.2.0.tar.gz, 10xGenomics) using Cell Ranger ATAC ver 1.2.0 (10x Genomics). The numbers of nucleus-hashing sequences mapped to each of the valid cells from Cell Ranger were counted using a Perl script, which is available as supplemental material.

### Single-cell ATAC-seq analysis

We re-aligned the obtained sequences to human GRCh38 (refdata-cellranger-atac-GRCh38-1.2.0, 10x Genomics) or mouse mm10 (refdata-cellranger-atac-mm10-1.2.0, 10x Genomics) references using Cell Ranger ATAC ver 1.2.0 (10x Genomics), separately. The quality of the libraries was assessed using ArchR [16]. The obtained peak counts were analyzed using Signac [17] and Seurat [22].

### Bulk ATAC-seq analysis

The bulk ATAC-seq libraries were analyzed, as we previously reported [23]. After assessing the qualities of the sequences using FastQC [24], the adapter sequences were trimmed with Cutadapt [25]. The adapter and quality trimmed sequences were aligned to mouse mm10 reference using BWA-mem software [26]. The peak-calling analysis on aligned reads was performed using MACS2 [27]. We calculated the reads in peak (RiP) with the ChIPQC Bioconductor package [28]. We used ChIP-R to identify the reproducible peaks [29]. We selected peaks shared across at least three biological replicates with the significance levels < 0.05 as high-confidence **<u>O</u>**pen **<u>C</u>**hromatin **<u>R</u>**egions (hc-OCRs). We used the hc-OCRs as well as entire OCRs to integrate OCRs we detected in snATAC-seq analyses.

### Identification of transcriptionally active putative enhancers

We combined all bulk ATAC-seq library aligned reads and performed peak-calling to generate a master OCR list for identification of enhancer regions and <u>T</u>ranscriptionally <u>A</u>ctive <u>P</u>utative <u>E</u>nhancers (TAPE), as previously reported [19]. We download and used the three HPC7 cell RNA-seq data from NCBI GEO website (GSE132724) [30]. The ATAC-seq results were merged and recentered using BEDTools (version2.28). All peaks smaller than 146 bp were removed to create a list of regions of open chromatin. Seqmonk (Babraham Institute) was used to identify intergenic regions of open chromatin (iROCs) and high-quality TAPEs by filtering probe list against known UCSC, Ensemble and Refseq genes curated lists for mm10. A final list of 145 high quality bidirectional TAPEs were identified. To map TAPEs to associated genes, transcription start sites located within 1Mb upstream or downstream from the center of the TAPE were identified. Then, Pearson’s correlations were calculated using counts for the TAPEs and associated genes.

### Assessing OCRs characteristics using publicly available datasets

To assess the characteristic of identified OCRs, we downloaded and used publicly available HPC7 datasets: a series of hematopoietic transcription factor ChIP-seq (GSE22178) [31] and CTCF ChIP-seq and Promoter capture Hi-C (pCHiC) (GSE129478) [32]. The overlap status was assessed using the findOverlapsOfPeaks function of the ChiPpeakAnno Bioconductor package [33]. The OCR annotation was performed using the annotatePeak function of the ChIPseeker Bioconductor package [34] with TxDb.Mmusculus.UCSC.mm10.knownGene annotation database [35]. Transcription factor binding motif enrichment analyses were performed using findMotifsGenome.pl from HOMER with mm10 reference [36]. All publicly available datasets were lifted to mm10 reference if the original analysis was performed on a different reference version, and MAC2 were used to call peaks on the bedGraph files.

## LIST OF ABBREVIATIONS

NuHash: Nucleus hashing
ATAC-seq: Assay for Transposase-Accessible Chromatin with high-throughput sequencing
ChIP-seq: chromatin immunoprecipitation followed by next generation sequencing
snATAC-seq: single-nuclear ATAC-seq
scRNA-seq: single-cell RNA-seq
UMI: unique molecular identifier sequences
GEM: Gel Beads-in-emulsion
TSS: transcription start site
OCRs: open-chromatin region
hc-OCRs: high-confidence Open Chromatin Regions
TADs: topologically associating domains
PIRs: promoter-interacting regions
FBS: fetal bovine serum
PBS: phosphate buffered saline
TAPE: transcriptionally active putative enhancers
pCHiC: promoter capture Hi-C

## DECLARATIONS

### Ethics approval and consent to participate

This study was approved by the Institutional Review Board (IRB) for Human Research at Albert Einstein College of Medicine, Bronx, NY (IRB Protocol# 2021-12969 and 2007-272).

### Consent for publication

All authors have read and approved the manuscript and gave their consent for submission and publication.

### Availability of data and materials

The sequencing data (bulk ATAC-seq and snATAC-seq) will be available at GEO before the publication (accession number pending). The detailed method of library preparation and the Perl script for demultiplexing NuHash used in this study are provided in the supplementary information.

### Authors’ contributions

Conceptualization, M.S.; methodology, B.S.B, H.N., D.R., S.B.M, and M.S.; formal analysis, T.V.T., E.S., R.A.D., D.Z., and M.S.; investigation, B.S.B. and M.S.; data curation, M.S.; writing— original draft preparation, M.S.; writing—review and editing, D.Z., B.E.M., J.M.G, and M.S.; visualization, M.S.; supervision, B.E.M. and J.M.G.; funding acquisition, M.S. All authors have read and agreed to the published version of the manuscript.

## Acknowledgements

The authors thank the continuous support from the Department of Genetics, Albert Einstein College of Medicine, for developing this method.

## Authors’ information

Current affiliations, Hiroko Nomaru, Thinkcyte Inc. Tokyo, Japan; Betelehem Solomon Bera, Center for Genetic Medicine, Children’s National Medical Center, Washington, DC, USA

## SUPPORTING INFORMATION CAPTIONS

**Supplementary file 1:** the detailed method for NuHash library preparation

**Supplementary file 2:** Supplementary figures

**Supplementary file 3:** NuHash demultiplexing script

**Supplementary file 4:** Supplementary tables

