## Supplementary file 1 for "An optimized approach for multiplexing single-nuclear ATAC-seq using oligonucleotide conjugated antibodies"

*Supplementary Methods*

### NuHash protocol

#### Step 1: Thawing frozen cells

This step is for the samples stored in a cell-freezing medium. If you need to store the samples, we recommend storing the cells with a cell-freezing medium (Cyrostor CS10 (StemCell technologies, cat# 07930) or CellBanker 2 (Amsbio, Cat # 11891)). To improve cell viability, freeze the cells slowly by reducing the temperature at approximately 1°C per minute using a controlled rate cryo-freezer or a cryo-freezing container.

If the starting material is a tissue, store the samples after the dissociation step.

**Materials needed at this step:**

Thawing medium: RPMI 1640 Medium (Gibco, cat# 11875093) supplemented 10% FBS (Gemini Bio, Cat# 100-106)

Resuspension buffer: 0.04% BSA (Sigma-Aldrich, cat# 126609-5GM) in PBS(-) (Gibco, cat# 10010031)

Resuspension buffer should be freshly prepared.

**Procedure:**

1. Thaw the cell in a 37 °C water bath until a tiny ice crystal remains (approximately 0.5 cm diameter)
2. Slowly transfer the cells into a 50 mL conical tube and sequentially dilute the cell suspension with a pre-warmed (37 °C) thawing medium.
   i. Add 1 mL of thawing medium to the tube drop by drop while gently agitating the tube (total volume, 2 ml)
   ii. Add 2 mL of thawing medium to the tube drop by drop while gently agitating the tube (total volume 4 ml)
   iii. Add 4 mL of thawing medium to the tube drop by drop while gently agitating the tube (total volume 8ml)
   iv. Add 8 mL of thawing medium to the tube drop by drop while gently agitating the tube (total volume 16 ml)
   v. Add 16 mL of thawing medium to the tube drop by drop while gently agitating the tube (total volume 32 ml)

3. Centrifuge at 300 RCF for 5 minutes at 4 °C.

**After this step, samples and medium/buffer should be kept on ice.**

4. Remove 31 mL of the supernatant without disrupting the cell pellet.
   We recommend doing two steps: 25 mL using a 25 ml disposable pipet, followed by 6 mL using a 10 mL disposable pipet.
5. Resuspend in the cell pellet in the remaining 1 mL of medium

6. Count the cell number using hemocytometer
7. Transfer the cell suspension to a 15 ml conical tube
8. Wash the 50 ml tube with 4 ml of thawing medium to collect remaining cells
9. Add additional 5 ml of thawing medium; total volume will be 10mL
10. Centrifuge at 300 RCF for 5 minutes at 4 °C
11. Remove supernatant without disturbing the pellet and resuspend in 1 ml of Resuspension buffer (combine tubes at this step, if it is applicable)
12. Gently mix with pipetting a few times
13. Transfer the cell suspension to a 2 ml tube
14. Wash 15 mL conical tube with 0.5 ml of Resuspension buffer for total volume of 1.5 mL

**Step 2: Nuclei isolation**

This step is for isolating nuclei. If you use fresh samples, start at this step.

**<u>Materials needed at this step:</u>**

| | |
|---|---|
| Wash buffer: | 10 mM Tris-HCl (pH 7.4), 10 mM NaCl, 3 mM $MgCl_2$, 1% BSA, 0.1% Tween-20 |
| Lysis buffer: | 10 mM Tris-HCl (pH 7.4), 10 mM NaCl, 3 mM $MgCl_2$, 0.1% Tween-20, 0.1% NP-40, 0.01% Digitonin (Invitrogen, cat#BN2006) |
| Staining buffer: | 2% BSA, 0.01% Tween-20 in PBS(-) |

All buffers should be freshly prepared and store them on ice.

**<u>Procedure</u>**

1. Centrifuge the 2 ml tube from the previous step at 300 RCF for 5 minutes at 4 °C
2. Remove supernatant without disturbing the pellet and transfer to 0.2 ml tube with total volume of 50 µl.
3. Centrifuge the 0.2 ml tube at 300 RCF for 5 minutes at 4 °C
4. Remove 45 µl of supernatant without disturbing the pellet
5. Add 45 µl of ice-cold lysis buffer to the tube and gently mix
6. Incubate the tube for 3 minutes on ice
7. Add 50 µl of ice-cold wash buffer (DO NOT mix)
8. Centrifuge at 500 RCF for 5 minutes at 4 °C
9. Remove 95 µl supernatant without disrupting the pellet.
10. Add 45 µl Staining buffer to the pellet (DO NOT mix)
11. Centrifuge at 500 RCF for 5 minutes at 4 °C
12. Carefully remove all supernatant using two steps, first using a 100 µl pipette (set at 40 µl) and then using a 10 µl pipette (set at 10 µL)
13. Resuspend the nuclei with 20 µl of ice-cold staining buffer
14. Count nuclei (take 2 µl of nuclei suspension and add 8 µl staining buffer)

**Step 3: Staining with nuclei-hashing antibody**

We stain the nuclei using the nuclei-hashing antibody (Nuc-Hash).

**Materials needed at this step:**

| | |
|---|---|
| Nuc-Hash antibody: | Custom oligo conjugated anti-Nuclear Pore Complex Proteins antibody (MAb414) |
| Fc Blocking reagent: | FcX (BioLegend, cat# 422301/ 101319) |
| Staining buffer: | 2% BSA, 0.01% Tween-20 in PBS(-) |

**Procedure**

1. Resuspend nuclei of each samples in 100 µl of staining buffer
2. Add 10 µl of Fc Blocking reagent
3. Incubate for 10 minutes on ice
4. Add NuHash antibody at 0.01 µg/50,000 nuclei ratio
5. Incubate for 20 minutes on ice
6. Add 1 ml of staining buffer and centrifuge at 500 RCF for 5 minutes at 4 °C
7. Remove supernatant and repeat step 6 twice
8. Carefully remove all supernatant and resuspend the nuclei with 5 µl of Diluted Nuclei buffer (10x Genomics, PN-20000153/20000207)
9. Count nuclei (take 1 µl of nuclei solution)
10. Combine equal number of nuclei per samples to make at 7700 nuclei/ µl solution

**After this step, follow the 10x scATAC library preparation procedure (from Step 1: Transposition)**
