## Supplementary file 2 for "An optimized approach for multiplexing single-nuclear ATAC-seq using oligonucleotide conjugated antibodies"

### Supplemental Figures

#### Supple Figure 1

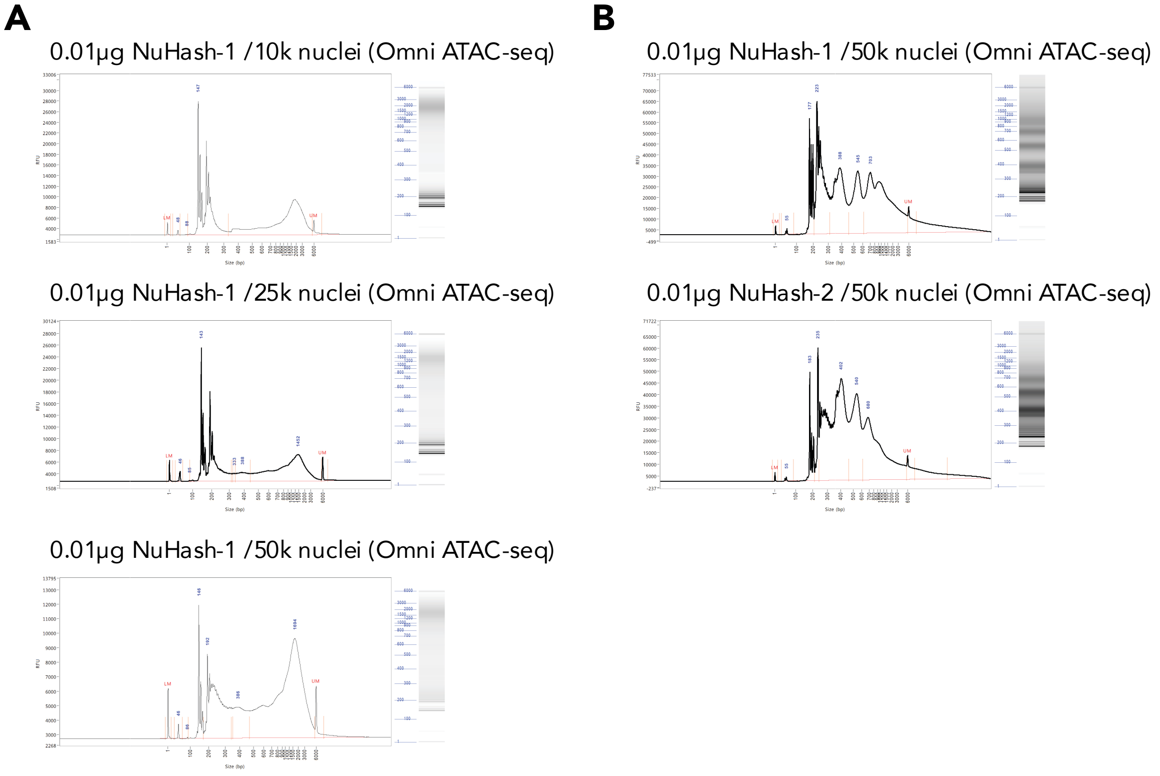


**Supple Figure 1: Optimizing NuHash antibody concentration to the number of nuclei.** We stained the human CD4+ T cell nuclei with different concentrations of NuHash antibody and generated bulk ATAC-seq (Omni- ATAC-seq) libraries to assess the proportions of NuHash and ATAC-seq products. (A) We stained with an antibody (NuHash-1) at three different concentrations. The panels showed the fragment distributions of each library before removing large fragments by size selection. (B) We stained with two different antibodies (NuHash-1 and NuHash-2)) at a 0.01 µg/50k nuclei concentration. The panel showed fragment distribution after removing large fragments by size selection.

#### Supple Figure 2

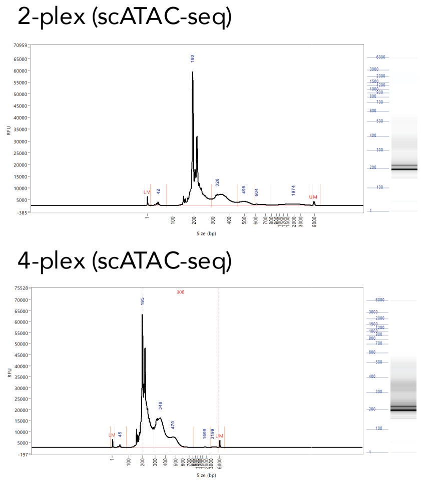


**Supple Figure 2: The library fragment distributions of NuHash scATAC-seq.**

We generated scATAC-seq libraries using multiplexing two (human and mouse samples, 2-plex) or four samples (two each human and mouse samples, 4-plex) using NuHash antibodies. The panels showed the fragment length distributions of the scATAC-seq library final products. The libraries contain the small fragments (NuHash products) and ATAC-seq banding pattern products.

#### Supple Figure 3

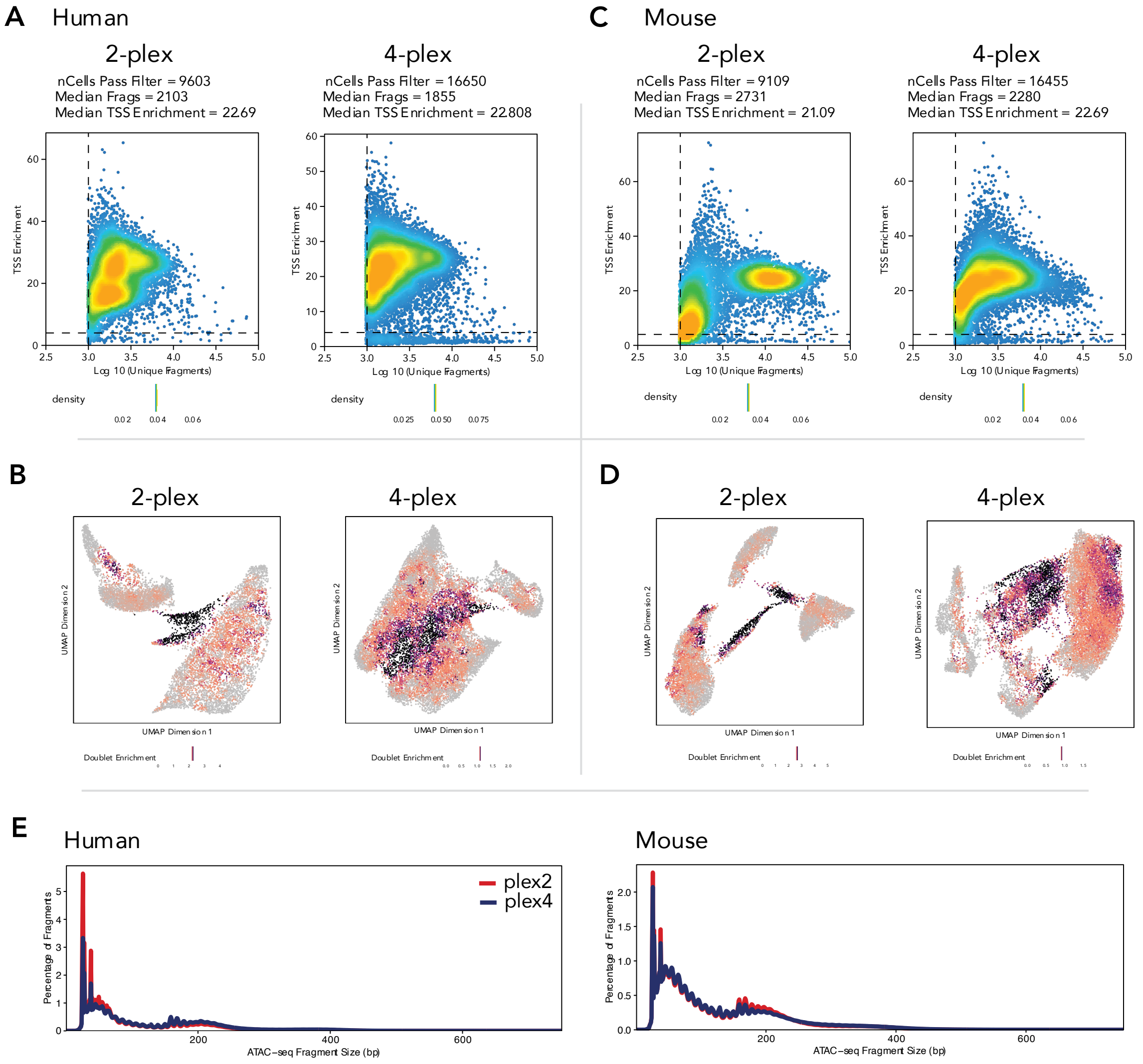


**Supple Figure 3: Quality assessment of the libraries.**

Transcription start site (TSS) enrichments scores and unique fragment numbers were plotted in **A** (human) and **C** (mouse) by the reference genomes. Colored nuclei with doublet enrichment scores were illustrated in **B** (human) and **D** (mouse). Panel **E** shows insert fragment length distributions.

#### Supple Figure 4

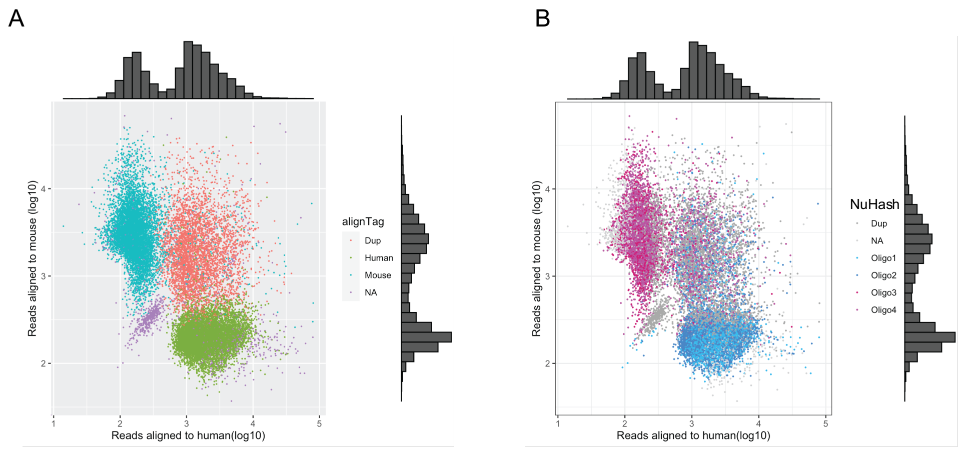


**Supple Figure 4: the fragment alignment status of each nucleus (4-plex).**

We plotted (**A**) the aligned read numbers to human or mouse references per nucleus, (**B**) the aligned read number per nucleus with colored by the NuHash count status.

#### Supple Figure 5

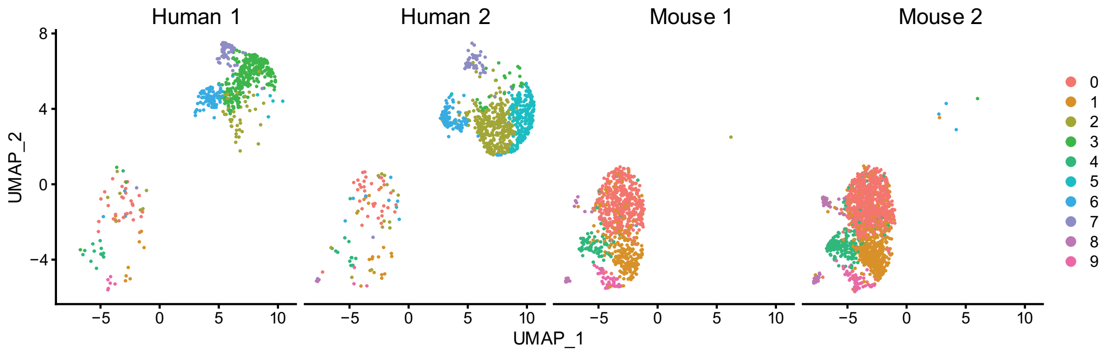


**Supple Figure 5: UMAP plots by samples**

UMAP plots showed a clear dissociation between human or mouse reference genome aligned nuclei.  The demultiplexed nuclei by the NuHash counting status were aligned to the reference genomes as expected.

#### Supple Figure 6

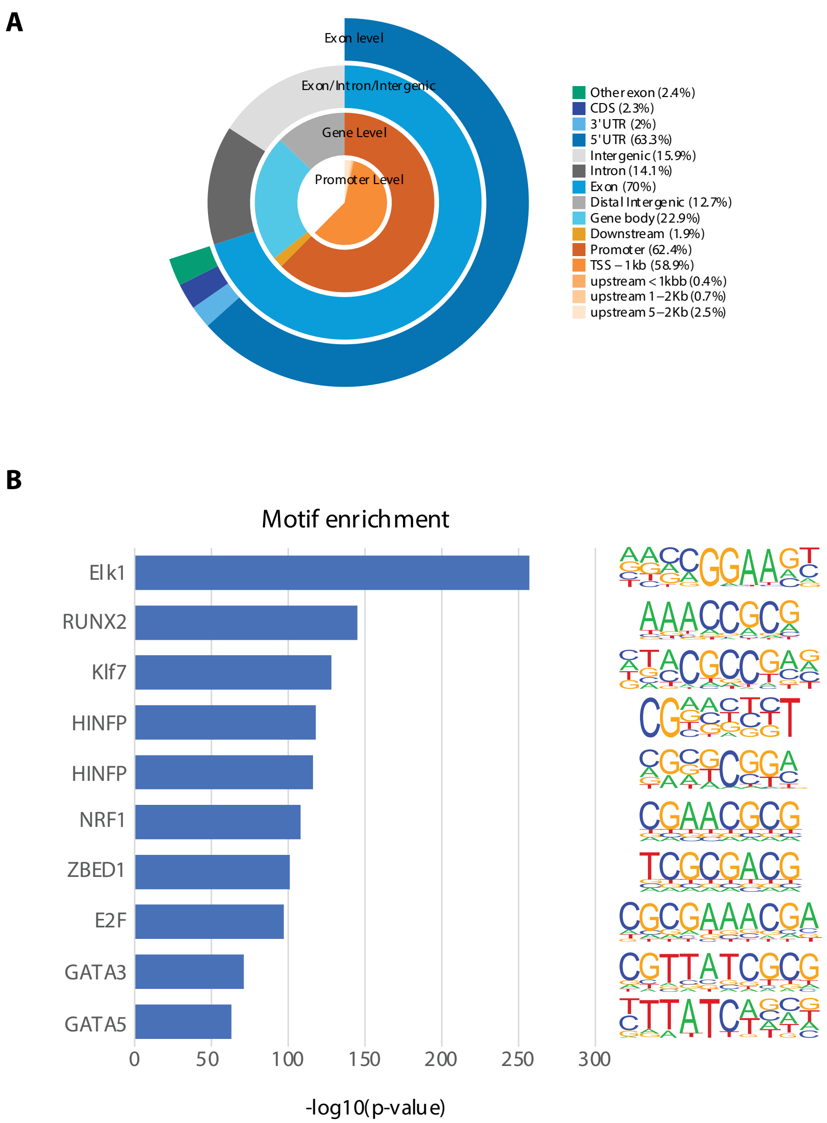


**Supple Figure 6: Peak characteristics of bulk ATAC-seq (HPC-7 cells)**

We plotted the peak characteristics of bulk ATAC-seq. Panel **A** shows distribution over different genomic features of the peak, and **B** indicates enriched transcription factor motifs in the detected peaks.
